# Integrative transcriptome analysis reveals distinct and common stress-responsive regulatory networks driving drought and heat responses in sorghum

**DOI:** 10.64898/2026.05.26.727923

**Authors:** Shikha Bharti, Nabanita Chattopadhyay, Salah E. Abdel-Ghany, Manohar Chakrabarti

## Abstract

Drought and heat are key abiotic stresses that severely affect crop production. Sorghum (*Sorghum bicolor L. Moench*), a drought-resilient cereal, serves as a model for studying abiotic stress responses in cereals. To elucidate the molecular responses to individual drought and heat stress, we conducted a transcriptome analysis of sorghum seedlings subjected to individual drought and heat treatment for 1h and 6h. Our analysis revealed distinct as well as overlapping patterns of gene expression between the two stresses at two time points, with a larger transcriptional shift observed at 6h of drought and heat treatment. We identified 410 and 4,136 differentially expressed genes (DEGs) in response to 1 and 6 h of drought treatments, respectively, whereas 1,807 and 2,776 DEGs were identified under 1 and 6 h of heat treatments. Among all four stress conditions, 32 common DEGs were identified. Genes encoding ion transporters were enriched among DEGs common in both 1h stress treatments. Genes involved in ribosome biogenesis were enriched among the common DEGs in both 6 h stress treatments. Specifically, drought-regulated genes were involved in ribosome biogenesis, whereas heat-induced genes were involved in protein folding and histone modifications. Enrichment of ribosome biogenesis in different sets of DEGs suggests that maintaining a balance between growth and survival by regulating protein synthesis may play a role in defining early stress response in sorghum. 6h of drought resulted in strong upregulation of abscisic acid and jasmonate-associated genes. Genes encoding bZIP, MYB, and HSF transcription factors displayed both stress-specific and common temporal regulation, suggesting vital regulatory roles of these transcription factors in mediating responses to drought and heat treatments. Extensive downregulation of genes encoding core histone proteins, in response to both 6h of drought and heat stress, was detected, indicating possible roles of chromatin structure and accessibility in mediating early responses to drought and heat treatments in sorghum.

## Introduction

Sorghum (*Sorghum bicolor* L. Moench) stands as a prominent and vital member of the Poaceae family and ranks fifth globally in both cultivated area and total production, thus outlining its significance in agriculture (FAOSTAT 2025). Sorghum has evolved and adapted to the unpredictable demands of semi-arid and dry habitats, displaying a number of uses including food, feed, and industrial raw materials (Silva et al. 2022). Millions of people in around 30 nations in subtropical and semi-arid Africa and Asia rely on sorghum as a staple food (FAOSTAT 2025). Beyond these areas, sorghum is also widely grown in the US, Latin America, and Australia, demonstrating its climate tolerance and global economic importance (Khalifa and Eltahir 2023). Many nations, particularly those in Africa and Asia, where sorghum grain is consumed as food, consider it a crop that ensures food security. High antioxidant and mineral contents, and a gluten-free label place sorghum as an excellent nutritionally-enriched food crop (Stamenković et al. 2020). Additionally, sorghum is gaining traction as a feed crop in high-input commercial farming; at the same time, its role in biofuel production continues to expand, driven by its versatility and resilience (Hariprasanna and Rakshit 2016). By integrating sorghum into biofuel production, agriculture can help address global energy challenges while reducing climate impact by 48% and fossil fuel depletion by 52%, as shown in a life cycle assessment of sweet sorghum biofuels in the U.S (Morrissey et al. 2021).

Sorghum uses the C4 photosynthesis pathway, which makes it more productive in hot and dry conditions than C3 crop plants (Tao et al. 2020). Furthermore, unlike maize (*Zea mays* L.), sorghum grows well on marginal lands with low fertility and high salt, showing its endurance beyond drought tolerance (Shan and Xu 2009). Sorghum thrives in semi-arid regions with optimal vegetative growth temperatures of 26–34°C daytime / 20–25°C nighttime (Prasad et al. 2015). However, global sorghum production occurs under unfavorable climatic conditions where seasonal temperatures regularly exceed 30°C. This includes countries, like Sudan, Mali, Niger, Senegal, and Saudi Arabia (Sultan et al. 2013). Recent climate analyses document extreme heat episodes well above 40°C throughout the sorghum-growing region in Sub-Saharan Africa, with Mali recording 48.5°C during the 2024 Sahel heatwave, Chad and Niger reaching 47–48°C, and South Sudan recording a temperature of 45°C in 2024 (Kunda et al. 2024). We know that global food security is gravely threatened by abiotic stresses, encompassing many factors, such as drought, salinity, heat, and cold, and are thus considered the main enemy of agricultural output. These stresses can act individually or in combination, further complicating crop resilience (Mukamuhirwa et al. 2019, Yanqing et al. 2026). It is even more critical to comprehend how these stresses impact stress-resilient crops, like sorghum, considering the impending threat of global climate change. In areas where food security is a problem, stresses like drought can not only lower grain yield but also worsen sorghum’s already poor protein digestibility. In regions where it is grown as a staple crop, this deterioration in quality might make malnutrition worse (Duodu et al. 2003). Plants rely on various physiological, morphological, and molecular defense mechanisms to withstand stress. However, the secondary effects of these stresses can be a lot more complex. These factors include the generation of reactive oxygen species (ROS), decreased metabolic activity, increased ion cytotoxicity, membrane instability, and increased cell death, all of which add another layer of difficulty to crop survival under such conditions (Zhu 2016; Saharan et al. 2022). Recent studies in wheat and rice corroborate these molecular signatures, emphasizing the role of ROS scavenging and membrane stability in stress resilience. For example, a transcriptomic meta-analysis in wheat identified differentially expressed genes linked to key pathways regulating ROS balance and membrane protection across multiple abiotic stresses (Ding et al. 2025). Similarly, in rice, genomic and transcriptomic approaches have highlighted pathways such as glutathione metabolism, peroxidase activation, and lipid membrane repair, which limit ROS accumulation and cellular damage under drought and heat stress (Ahmad 2022). These findings indicate that ROS detoxification and membrane stabilization represent conserved and critical mechanisms in major cereals for combating abiotic stresses.

Thus, the importance of sorghum in maintaining food security and sustainable agriculture amid rapidly changing climate conditions is paramount. Sorghum is the perfect model for analyzing the molecular processes involved in drought and heat tolerance due to its resilience to these abiotic stresses. Prior transcriptome studies either looked at responses to drought stress in a particular sorghum cultivar or in two contrasting sorghum cultivars (Abdel-Ghany et al. 2020, Azzouz-Olden et al. 2020, Wang et al. 2025). A recent study explored responses of sorghum seedlings to 1, 2, 5, and 7 days of PEG-simulated drought stress (Ma et al. 2026). There is a need to carry out comparative analysis to decipher the uniqueness and overlap among molecular responses to different stresses in sorghum. Under this backdrop, using a genome-wide 3′-end sequencing approach (Chakrabarti et al. 2018), the current study aims to identify key genes involved in regulating early responses to individual drought and heat treatments at two time points, including 1 and 6 hours. Our transcriptome analysis elucidated time-dependent changes in gene expression profiles in responses to individual drought and heat stress and deciphered commonalities and uniqueness between transcriptional responses to these individual stress treatments in sorghum.

## Materials and methods

### Experimental design, tissue collection, and 3′-end sequencing

An abiotic stress experiment was conducted with the *Sorghum bicolor* genotype ‘BTx623’, an early-maturing, short-statured, inbred variety, with a well-annotated reference genome, as described before (Abdel-Ghany et al. 2016). Seeds were surface sterilized using a 20 % bleach solution for half an hour, and they were allowed to germinate for 24 hours on moist filter paper. After germinating, the seedlings were placed in culture tubes and were grown on 0.5x Hoagland nutrient solution as described previously for an additional 7 days in a growth chamber (Abdel-Ghany et al. 2016). Our experimental setup includes two treatments (individual drought and heat) and one control (the same control was used for the drought and heat treatments), and two time points (1 and 6h). On the eighth day, the nutrient medium was replaced with 5 ml of 0.5x Hoagland solution for control samples. The nutrient medium was replaced with 5 ml of 0.5x Hoagland solution containing 20% PEG-8000 for drought-treated samples. For the heat treatment, sorghum seedlings were grown in 5 ml of 0.5x Hoagland solution in a growth chamber whose temperature was set at 45°C. Control and drought-treated samples were kept in a growth chamber set at 26°C. After being treated for one and six hours, tissue samples were collected, flash frozen in liquid nitrogen, and total RNA was extracted with the miRNeasy Mini Kit along with on-column DNAse I digestion (QIAGEN) following the manufacturer’s protocol. The experiment was conducted with three biological replicates. The 3′-end sequencing libraries were prepared and sequenced on the Illumina platform as part of our prior study (Chakrabarti et al. 2020). The 3′-end sequencing data was deposited in the NCBI-SRA (Sequence Read Archive) under the BioProject accession number PRJNA523821.

### Processing of 3′-end sequencing data

The sequence files of each replicate of drought-treated and heat-treated samples at 1 and 6h after the stress treatments and their corresponding control samples were retrieved from NCBI SRA. In total, there were 18 samples. Processing of raw 3′-end sequencing reads, and transcriptomic analysis were performed using various tools available in CLC Genomics Workbench (v 23.0.4) as described previously (Chakrabarti et al. 2020). Raw 3′-end sequencing reads were trimmed using the ’Trim Reads’ tool to remove Illumina adapter sequences and oligo-dT sequences. Trimmed reads were mapped to ribosomal RNA genes, mitochondrial, and chloroplast genomes using the ‘Map reads to Reference’ tool using the following mapping parameters: match score-1, mismatch cost-2, insertion cost-3, deletion cost-3, length fraction-0.5, and similarity fraction-0.6. Unmapped reads were collected and were aligned to the *Sorghum bicolor* reference genome [version 3.1.1, downloaded from Phytozome (http://phytozome.jgi.doe.gov)] using the ‘RNA-seq Analysis’ tool in the CLC Genomics Workbench (version 23.0.4). For the mapping to the sorghum genome, the following parameters were used: match score-1, mismatch cost-2, insertion cost-3, deletion cost-3, length fraction-0.6, similarity fraction-0.8, and maximum number of hits for a read-10.

### Differential gene expression analysis

Differential gene expression analysis was conducted using the CLC Genomics Workbench (version 23.0.4). Mapped read value for each gene was transformed and normalized to ensure comparability of gene expression values among replicates and among samples. Expression counts were transformed by adding 1. Normalization was conducted using the counts per million (CPM) method, where mapped read counts were divided by the number of reads in a particular library and then multiplied by 1,000,000. Low-expressed genes were excluded from the subsequent analysis, and genes with > 2 CPM (average of three replicates of a treatment) in at least one stress-treated or the respective control sample were retained for downstream analyses. Baggerley’s test was conducted to identify differentially expressed genes (DEGs). Genes that displayed at least a 2-fold change (based on CPM values, average of three replicates) and an FDR (False Discovery Rate)-corrected p-value of <0.05 in a comparison between stress-treated samples and corresponding control samples were designated as DEGs. Expressions of ten stress-responsive DEGs were validated using an independent RNAseq dataset (NCBI SRA BioProject accession: PRJNA523862). The RNAseq dataset was generated using the same BTx623 sorghum genotype, and drought and heat treatments were performed in the same way as mentioned in the preceding section. The RNAseq data was analyzed in the same way as the 3’-end sequencing dataset. For the RNAseq dataset, genes that displayed at least a 2-fold change in the normalized expression (average of three replicates) and a p-value of <0.05 in a comparison between stress-treated and control samples were considered as DEGs. Fold change values (Stress divided by control, based on CPM values of three biological replicates) of these ten DEGs from the 3 -end sequencing and RNAseq datasets were compared.

### Gene Ontology analysis

For the Gene Ontology (GO) analysis, we employed the AgriGO v2.0 web-based platform (Tian et al. 2017). GO analysis was done using ‘Singular Enrichment Analysis’ (SEA), specifically utilizing the "Complete GO". We used a customized background, which retained all genes with expression >2 CPM (counts per million) in at least one treatment (an average of three replicates), which includes GO annotations based on *Sorghum bicolor* genome version 3.1.1 (Phytozome v11.0). To ensure statistical significance, we applied the Hochberg (FDR) test for multiple test adjustments, with a significance level of 0.05. Additionally, we set the minimum number of mapping entries to 3. To visualize enriched GO terms and pathways more effectively, we utilized REVIGO with default parameters (Supek et al. 2011). REVIGO summarizes and visualizes GO terms by consolidating redundant terms, thus providing a clearer overview of the enriched GO categories.

### Pathway enrichment analysis using PageMan and MapMan

To visualize the under- and over-representation of different biological pathways, PageMan, a plugin of MapMan, was utilized (Usadel et al. 2006), which provided a broad overview of biological processes impacted in responses to drought and heat stress in sorghum. The over- and under-representation analyses in PageMan were carried out using the ‘ORA_Fisher’ test with the Benjamini-Hochberg multiple testing correction and an ORA cutoff value of 1. Pathways related to lipid metabolism, hormone metabolism, cell wall biosynthesis and degradation, photosynthesis, redox regulation, amino acid metabolism, responses to abiotic stresses, and transcriptional regulators were visualized (**Figure S1**).

We further used MapMan (Usadel et al. 2009) to visualize different functional categories of genes that were significantly altered in responses to 1 and 6h of individual drought and heat treatments as compared to their respective control samples in sorghum. Four pair-wise comparisons were conducted, which include comparison between 1h drought-treated (D1_C1), 6h drought-treated (D6_C6), 1h heat-treated (H1_C1), and 6h heat-treated (H6_C6) samples with their corresponding control samples (C1 refers to control samples for both 1h drought- and heat-treated samples, whereas C6 refers to control samples for 6h drought- and heat-treated samples). MapMan allowed us to categorize DEGs into specific functional bins and identify significant changes in gene expression associated with these stress treatments.

## Results

### Drought and heat treatments drastically alter gene expression patterns in sorghum seedlings

Seedlings of the sorghum cultivar BTx623 were chosen to study responses to individual drought and heat treatments. In total, Illumina sequencing data for 18 libraries, consisting of three replicates of drought- and heat-treated samples and their corresponding control sample, were used for this study. In this study, 3′-end sequencing reads were used to measure global gene expression, as done by previous studies (De Lorenzo et al. 2017, Chakrabarti et al. 2018). A summary of reads retained after each step during data processing is represented in Supplemental **Table S1**. We conducted differential gene expression analyses between stress-treated and corresponding control samples at 1 and 6h after the stress treatment (**Fig.1A**). In total, 6,773 DEGs were identified between stress-treated and control samples at two time points (**Fig.1B**). Considerable changes in gene expression were observed in response to all stress conditions, except in response to 1h of drought treatment (**Fig.1B**).

**FIGURE 1.**
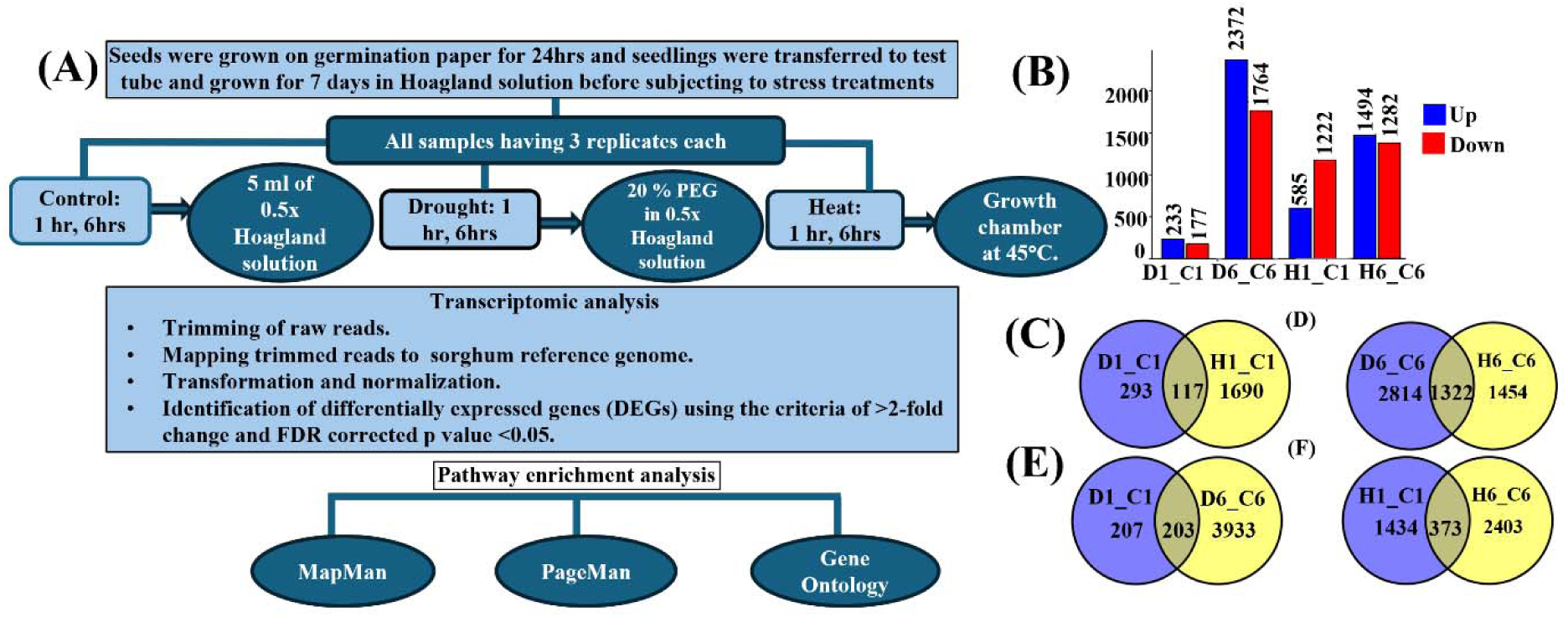
Overview of experimental design and differential gene expression analysis in sorghum seedlings in responses to 1h and 6h of individual drought and heat treatment. (A) Schematic representation of the experimental design. (B) Bar graph showing the number of up- and down-regulated differentially expressed genes (DEGs) across different stress treatments. (C)–(F) Venn diagrams showing unique and shared DEGs in seedlings subjected to 1h and 6h of drought and heat treatment. D1, H1, and C1 represent drought, heat, and control samples at 1h, while D6, H6, and C6 correspond to the same samples at 6h. D1_C1, H1_C1, D6_C6, and H6_C6 represent sets of DEGs identified in responses to 1h drought, 1h heat, 6h drought, and 6h heat treatment, respectively. C1 represents control samples for both 1h drought- and heat-treated samples. C6 represents control samples for both 6h drought- and heat-treated samples.

### Transcriptional responses of sorghum seedlings to drought and heat treatments at two different time points

We identified 410 DEGs after 1h of drought (D1) treatment as compared to the control condition, where 233 and 177 DEGs were up- and down-regulated in drought-treated samples as compared to control samples (**Fig.1B**). In contrast, 6h of drought treatment (D6) resulted in 4,136 DEGs, and out of those, 2,372 and 1,764 DEGs were found to be up- and down-regulated in drought-treated samples as compared to control samples (**Fig.1B**). 1h of heat stress exerted a more significant effect as compared to 1h of drought stress. For 1h of heat stress (H1), we had 1807 genes showing differential expression, and out of these, 585 genes were up-regulated, and 1222 genes were down-regulated. For 6h heat stress (H6), 2776 genes were differentially expressed, with 1494 and 1282 genes showing up- and down-regulation, respectively, in response to heat treatment (**Fig.1B**). Temporal analysis revealed significant variation between responses to 1 and 6h of drought treatment. Our study identified 207 and 3933 DEGs that were unique to 1 and 6h of drought treatments, respectively (**Fig.1E**). Whereas 203 genes were differentially expressed in responses to both 1 and 6h of drought treatments. For heat stress, 1434 and 2403 genes were specifically differentially expressed in responses to 1 and 6h of heat treatments, respectively, while 373 genes were differentially expressed in responses to both 1 and 6h of heat treatments (**Fig.1F**).

GO analysis was conducted to elucidate enrichment of genes involved in specific molecular processes in responses to drought and heat treatments in sorghum seedlings. DEGs identified after 6h of drought treatment displayed enrichment of genes that are involved in ribosomal biogenesis, and that encode oxidoreductases and hydrolases (**Fig.S2A**). GO analysis of the 203 DEGs common in 1 and 6h of drought treatment (common in D1_C1 and D6_C6) exhibited enrichment of genes that encode transcription factors, phosphoprotein phosphatases, and that possess hydrolase activity (**Fig.2A**). GO analysis of a set of DEGs specific to the 1h of drought treatment (specific in D1_C1) didn’t reveal enrichment of any specific class of genes. Similarly, GO analysis of the 3933 DEGs specific for 6h of drought treatment (specific in D6_C6) also revealed enrichment of genes implicated in maintaining the structural integrity of ribosomes (**Fig.2A**).

**FIGURE 2.**
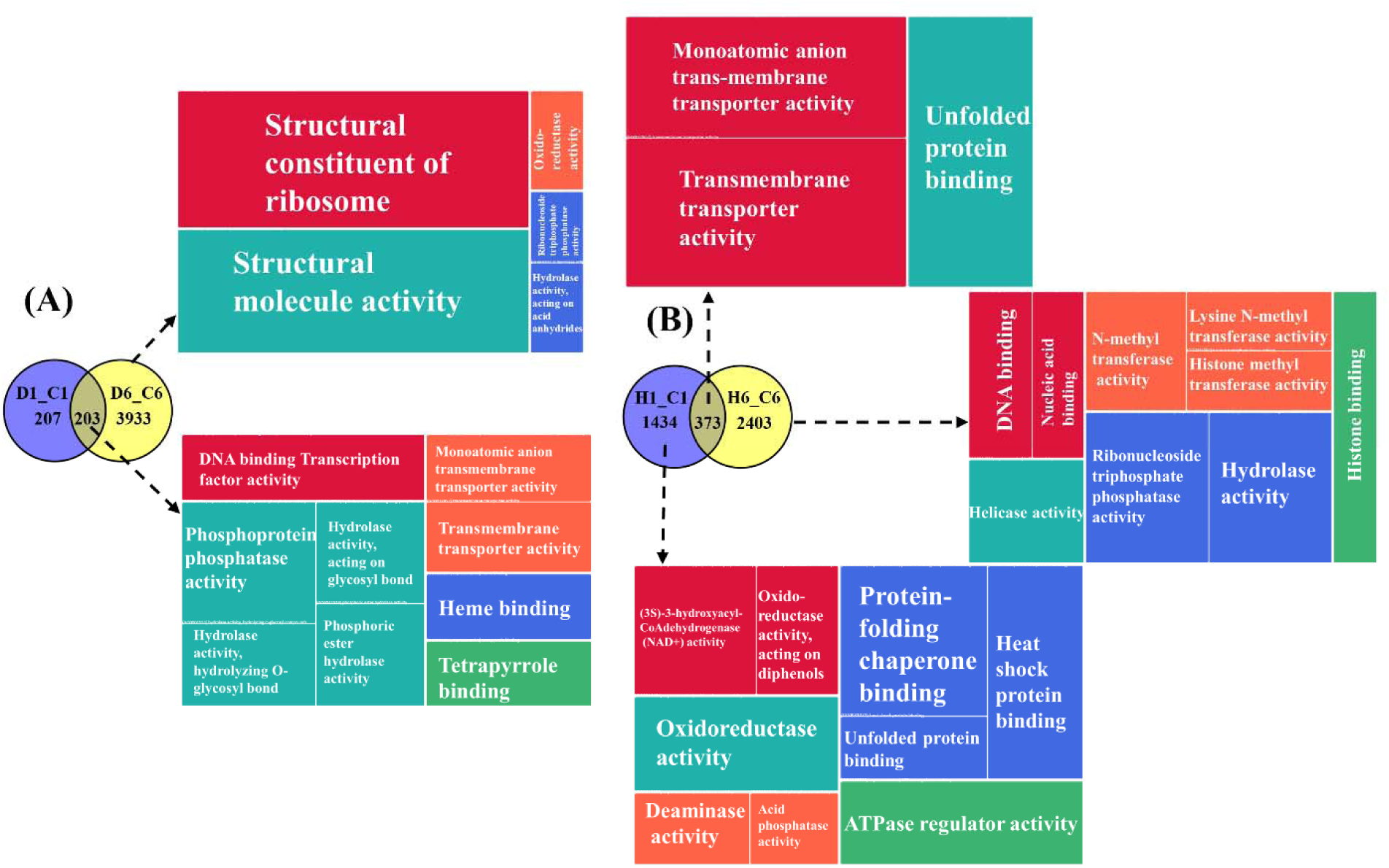
Comparative GO analyses of DEGs in response to 1 and 6h of drought (D1_C1 & D6_C6) and 1 and 6h of heat treatment (H1_C1 & H6_C6), respectively. (A) and. **(B)** represent GO categories enriched in sets of DEGs that were either specific or overlapping in responses to 1h and 6h of drought, and heat treatment, respectively. The accompanying TreeMaps display enriched GO terms under the ‘Molecular Function’ category. No significant molecular function terms were enriched for DEGs that were specific to 1h drought stress (D1_C1). D1, H1, and C1 represent drought, heat, and control samples at 1h. D6, H6, and C6 represent the same samples at 6h. D1_C1, H1_C1, D6_C6, and H6_C6 correspond to sets of DEGs identified in responses to 1h drought, 1h heat, 6h drought, and 6h heat treatment, respectively.

In contrast, DEGs after 6h heat treatment were enriched for genes that encode hydrolases and that are involved in the regulation of ribonucleoside triphosphate activity (**Fig. S2B**). GO analysis of 373 DEGs common in 1 and 6h of heat treatment (common in H1_C1 and H6_C6) revealed enrichment of genes involved in anion transport activity, protein folding, and transmembrane transporter activity. (**Fig.2B**). A set of 1,434 DEGs specific to the 1h heat treatment (H1_C1) exhibited enrichment of genes involved in protein folding, ATPase activity, and genes that encode heat shock proteins (HSPs) and oxidoreductases, such as 3-hydroxyacyl-CoA dehydrogenase. A set of 2,403 DEGs specific for 6h of heat treatment (specific in H6_C6) revealed enrichment of genes that are implicated in histone binding, ribonucleoside triphosphate phosphatase activity, and that encode hydrolases (**Fig.2B**). These results suggest that heat stress triggered immediate changes in ribonucleoside triphosphate phosphatase activity implicated in cell signaling and cellular transport. Heat stress also led to alterations in chromatin accessibility and protein folding.

### Commonalities and uniqueness in responses to drought and heat treatment

We conducted a comparative study to elucidate common and unique transcriptional responses to individual drought and heat treatments (**Fig.1C and D**). Our analysis identified that 293 and 1,690 genes were differentially expressed specifically when subjected to 1h of drought and heat treatment, respectively. Notably, 117 genes were differentially expressed in responses to both 1h of individual drought and heat treatment (**Fig.1C**). Similarly, for 6h drought and heat treatment, 2,814 and 1,454 genes showed differential expression specifically after 6h of drought and heat treatment, respectively, and 1,322 genes showed differential expression in responses to 6h treatment of both stresses (**Fig.1D**).

A set of 117 DEGs that were differentially expressed both after 1h of drought and heat treatments (common in D1_C1 and H1_C1), displayed enrichment of genes that encode transmembrane ion transporters (**Fig.3A**). A set of 293 DEGs specific for 1h of drought stress (specific in D1_C1) revealed enrichment of genes that encode protein phosphatases, transcription factors, and oxidoreductases. A set of 1,690 DEGs specific for 1h of heat stress (specific in H1_C1) exhibited enrichment of genes that encode oxidoreductases, heat shock proteins, and that are implicated in protein folding (**Fig.3A**). A set of 1,322 DEGs that were differentially expressed both after 6h of drought and heat treatments (common in D6_C6 and H6_C6) showed enrichment of genes implicated in ribosome biogenesis. Similarly, 2,814 DEGs specific for 6h of drought treatment (specific in D6_C6) exhibited enrichment of genes implicated in ribosome biogenesis. A set of 1,454 DEGs specific to 6h of heat treatment (H6_C6) displayed enrichment of terms related to histone methyltransferase activities (**Fig.3B**). Altogether, our results revealed involvement of ion transporters in regulating responses to both1h stress treatments. Transcription factors and post-translational protein modifications emerged as vital regulators of response to 1h drought treatment, whereas ATPase involved in active transport and protein folding appeared to regulate response to 1h heat treatment. Our results also identified biogenesis of ribosomes, which is crucial for protein synthesis, as a key regulator of response to 6h drought treatment. Histone modification emerged as a possible regulator of response to 6h of heat treatment.

**FIGURE 3.**
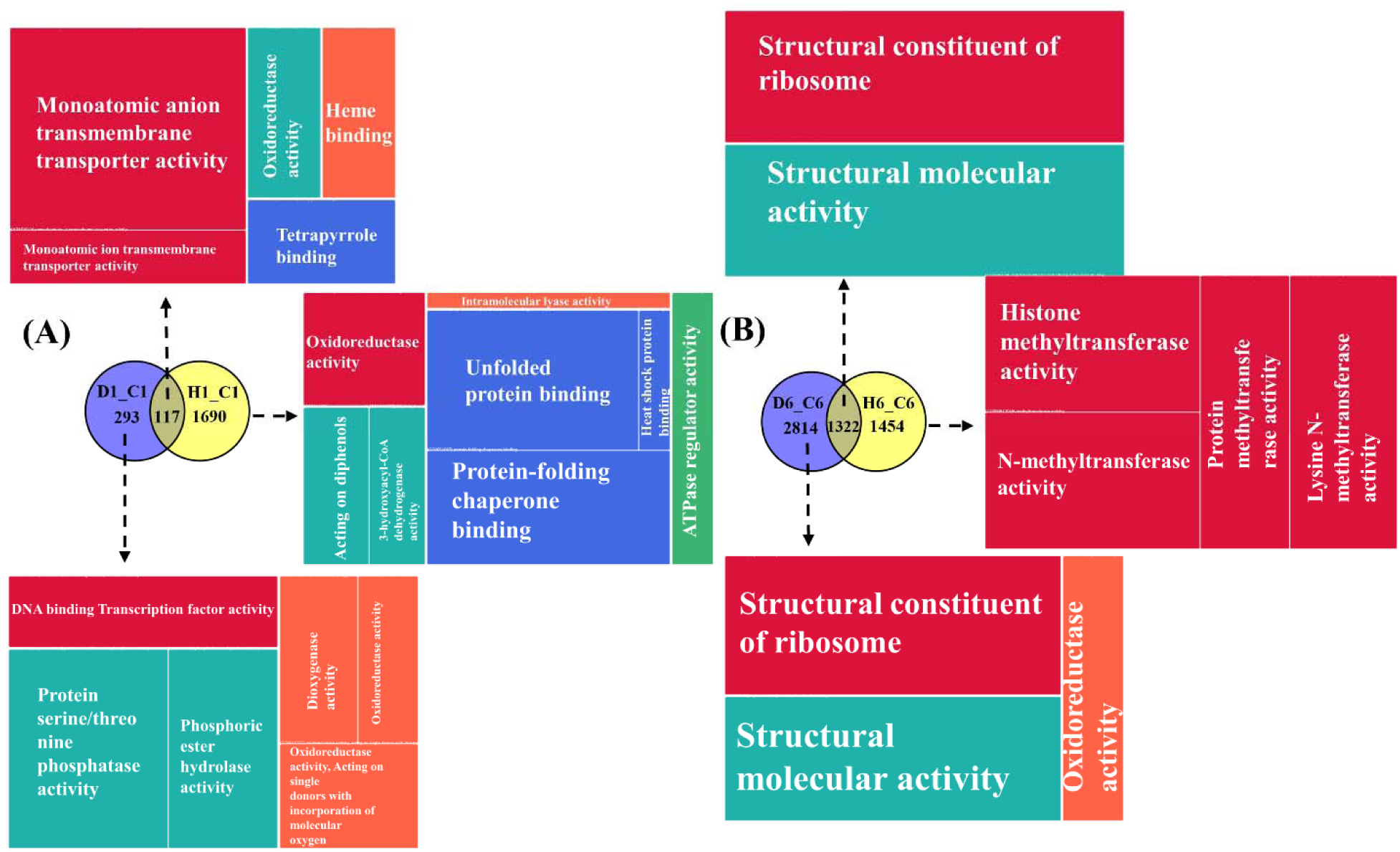
Comparative GO analyses of DEGs in responses to drought and heat treatments at 1 and 6h. (A) and. **(B)** represent GO categories enriched in sets of DEGs that were either specific or overlapping in responses to 1h of drought (D1_C1) and heat (H1_C1), and 6h of drought (D6_C6) and heat treatment (H6_C6), respectively. D1, H1, and C1 correspond to drought, heat, and control samples at 1h, while D6, H6, and C6 refer to the same samples at 6h. D1_C1, H1_C1, D6_C6, and H6_C6 represent sets of DEGs identified in responses to 1h drought, 1h heat, 6h drought, and 6h heat treatment, respectively.

We identified 32 genes that were differentially expressed in responses to all 1 and 6h of abiotic stress treatments as compared to corresponding control samples. Among these, 6 genes exhibited up-regulation, while 26 genes were down-regulated in responses to all stress treatments (**Fig.4**). The set of up-regulated genes was implicated in salicylic acid metabolism and mediating responses to heat shock, whereas the group of down-regulated genes encodes different ion transporters, ribosomal proteins, oxidoreductases, and others (**Fig.4**).

**FIGURE 4.**
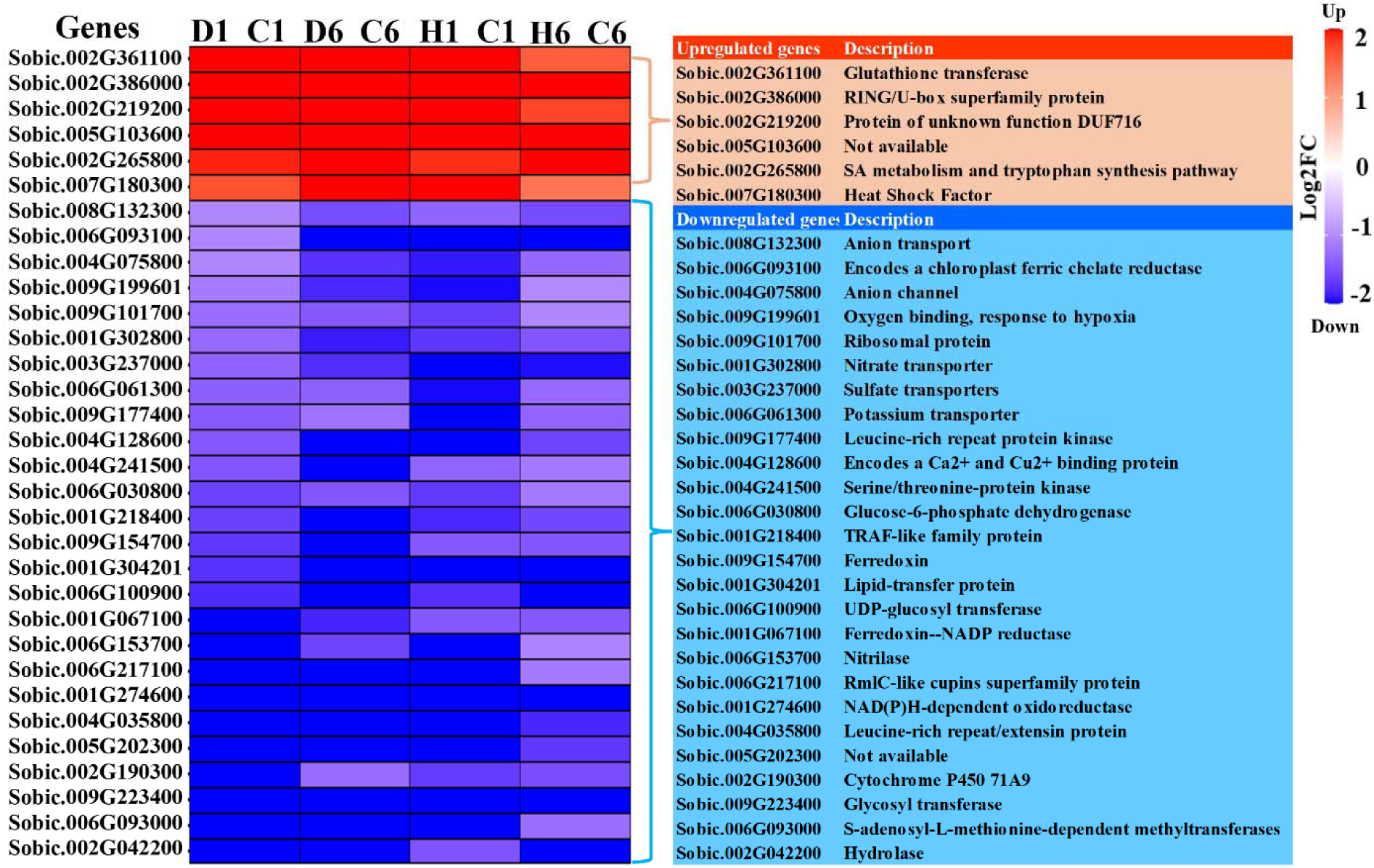
Heatmap showing the expressed across four stress conditions. D1, H1, and C1 refer to drought, heat, and control samples at 1h. D6, H6, and C6 refer to the same samples at 6h. D1_C1, D6_C6, H1_C1, and H6_C6 represent comparisons of 1h drought-treated, 6h drought-treated, 1h heat-treated, and 6h heat-treated samples with their corresponding control samples, respectively. Gene expression levels are represented as Log_2_ Fold Change (normalized expression in stress-treated sample/normalized expression in control sample), with red and blue indicating up- and down-regulation, respectively.

### Effects of drought and heat treatments on phytohormone and secondary metabolism

To assess and visualize an overview of drought- and heat-induced changes in gene expression in multiple stress conditions, the PageMan tool was used (Usadel et al. 2006). Our analysis revealed drought- and heat-induced changes in the expressions of genes involved in phytohormone-related processes and in secondary metabolism (**Fig.S1**). We further mapped and visualized stress-induced DEGs into different MapMan bins. We did not notice statistically significant enrichment of genes involved in ABA and JA-related processes, except in the up-regulated DEGs after 1h drought treatment in our PageMan analysis. However, since two phytohormones, ABA and JA, are widely regarded as major regulators of stress response in different plant species (Khan 2025, Li et al. 2025), we assessed expression of genes involved in ABA- and JA-related processes in all four stress conditions. We identified 37 genes involved in ABA-related processes (e.g., synthesis, signaling, and response) that exhibited differential expression in at least one stress condition (D1, D6, H1, and H6) as compared to their corresponding control condition (**Fig.S3A & S4A**). Two genes, including *Sobic.002G107000* (encodes an aldehyde oxidase) and *Sobic.004G268700* (encodes ABA 8’-hydroxylase), were differentially expressed only in response to 1h of heat treatment (**Fig.S3A**). 17 genes were differentially expressed exclusively in response to 6h of drought treatment (**Fig.S3A**). This set of genes included *Sobic.001G155300* and *Sobic.002g037400* (both encode 9-cis-epoxycarotenoid dioxygenase, involved in ABA biosynthesis), *Sobic.002G360701* (encodes protein phosphatase 2C), *Sobic.004G309600* [encodes an ABRE (ABA-responsive element)-binding protein], and others. Four genes were differentially expressed specifically in response to 6h of heat treatment, and an additional five genes were differentially expressed in response to both 6h of drought and heat treatment (**Fig.S3A**).

Other than ABA, drought and heat treatments altered expressions of jasmonate (JA)-related genes as compared to control conditions. We identified 21 JA-related genes that were differentially expressed in response to at least one stress treatment. 1h of drought treatment resulted in the up-regulation of three and down-regulation of one JA-related gene. In comparison, 6h of drought treatment resulted in up-regulation of 13 and down-regulation of two JA-related genes (**Fig.S3B & S4B**). JA-related genes also displayed differential expression in response to heat treatments. 1h of heat treatment resulted in up- and down-regulation of two and three JA-related genes, respectively. On the contrary, 6h of heat treatment resulted in up- and down-regulation of two JA-related genes each (**Table S2**). Two JA-related genes were differentially expressed in response to both 6h stress treatments (**Fig.S3B**). Altogether, our study identified sets of ABA and JA-related genes that were either differentially expressed uniquely in response to drought or heat stress, or in response to both individual stress treatments.

Secondary metabolism was not significantly affected by 1h of drought treatment, but it was significantly affected by 1h of heat treatment (**Fig.S5A & C**). After 1h heat treatment, genes related to lignins, other phenylpropanoids, flavanols, dihydroflavonols, anthocyanins, and glucosinolates were mostly down-regulated (**Fig.S5C**). In contrast, the 6h of drought and heat treatments did not induce significant down-regulation of genes associated with secondary metabolism-related pathways. In response to 6h of drought treatment, genes related to terpenoids, anthocyanins, and flavonols were mostly up-regulated, and genes related to glucosinolates were primarily down-regulated, while genes related to lignins, other phenylpropanoids, and dihydroflavonols were both up- and down-regulated. In the case of 6h of heat treatment, genes related to terpenoids were up-regulated, genes associated with lignins were primarily down-regulated, whereas genes related to phenylpropanoids showed both up- and down-regulation (**Fig.S5B & D**).

### Drought and heat stress altered the expression of genes encoding transcription factors

Our MapMan analysis revealed the temporal effects of drought and heat stress on the expression of genes encoding different transcription factors (**Fig.S6**). Our MapMan analysis showed that at 1h of drought, genes encoding bZIP and MYB transcription factors displayed minimal expression changes; however, by 6h, these genes showed substantial expression changes (**Fig.S6**). Hence, genes encoding bZIP and MYB transcription factors were studied further. A total of 32 genes encoding bZIP transcription factors were differentially expressed in response to at least one stress condition (**Fig.S3C & Fig.5A**). 1h of drought treatment resulted in up- and down-regulation of three genes and one gene that encodes bZIP transcription factors. Similarly, 1h of heat treatment resulted in up-regulation of two and down-regulation of four genes that encode bZIP transcription factors. In total, 17 bZIP-encoding genes were up-regulated, while five were down-regulated in response to 6h of drought treatment. In comparison, 6h of heat stress resulted in the up-regulation of six and down-regulation of five bZIP-encoding genes (**Table S2**). In total, one, 14, four, and four bZIP-encoding genes were exclusively differentially expressed in responses to 1h of drought, 6h of drought, 1h of heat, and 6h of heat treatment, respectively (**Fig.S3C**). Six bZIP-encoding genes that were commonly differentially expressed in response to both 6h of drought and heat treatment included *Sobic.010G194900* (a likely ortholog of arabidopsis bZIP60, associated with the endoplasmic reticulum stress response) (**Fig.5A**).

**Figure 5.**
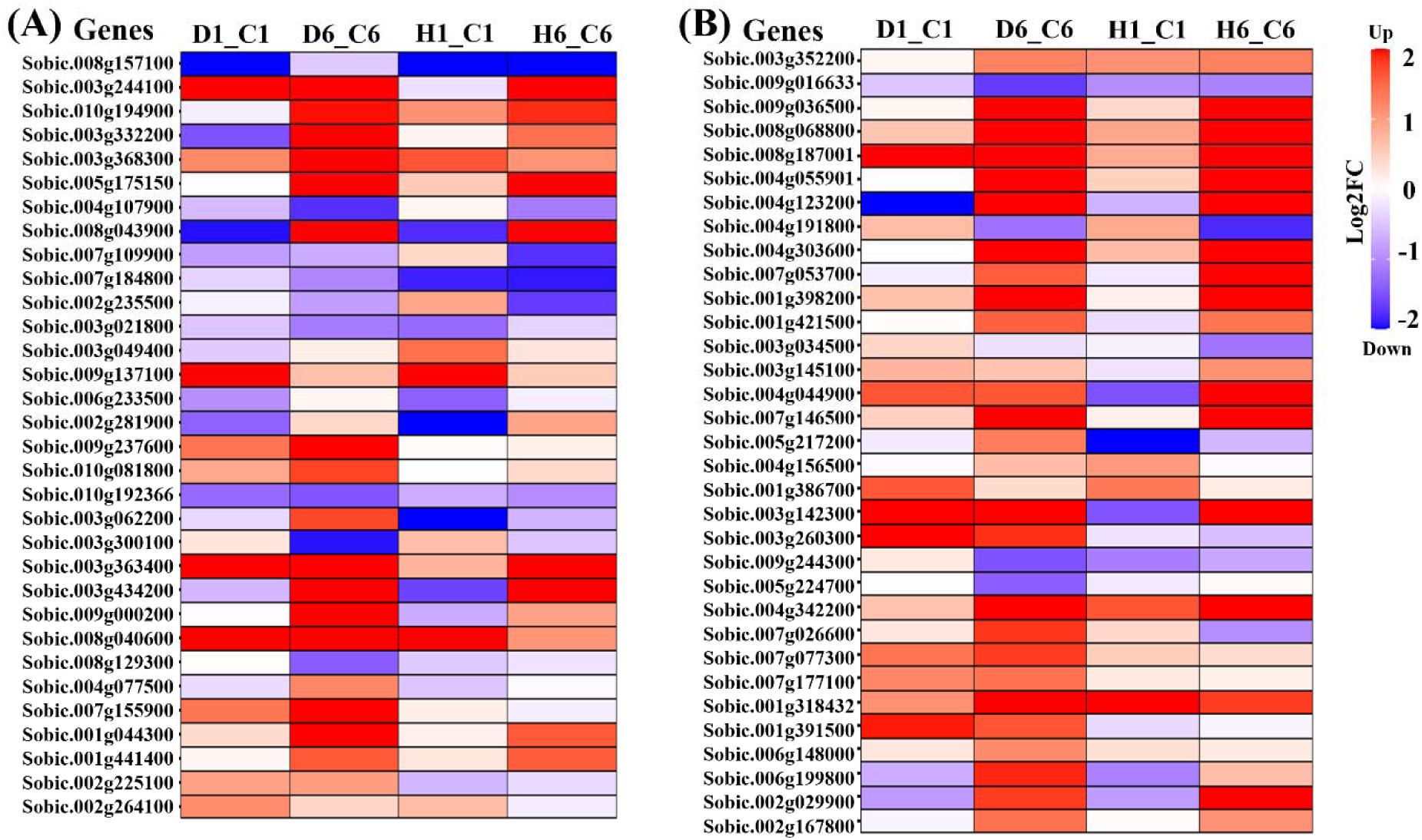
Heatmaps showing DEGs encoding bZIP and MYB transcription factors in responses to 1 and 6h of drought and heat treatments. D1_C1, D6_C6, H1_C1, and H6_C6 represent comparisons of 1h drought-treated, 6h drought-treated, 1h heat-treated, and 6h heat-treated samples with their corresponding control samples, respectively. **(A)** and **(B)** represent heatmaps showing the expression patterns of genes encoding bZIP and MYB transcription factors, respectively. Gene expression values are represented as Log_2_ Fold Change (normalized expression in stress-treated sample/normalized expression in control sample), with red and blue indicating up- and down-regulation, respectively.

The current study also identified a total of 33 MYB transcription factor-encoding genes that were differentially expressed genes (DEGs) in response to at least one stress treatment (**Fig.S3D** & **Fig.5B**). 1h of heat treatment resulted in up- and down-regulation of two and one MYB transcription factor-encoding genes, respectively. No MYB transcription factor-encoding gene showed differential expression in response to 1h of drought treatment. 6h of drought treatment resulted in up- and down-regulation of 22 and four MYB transcription factor-encoding genes, respectively, whereas 6h of heat treatment resulted in up- and down-regulation of 13 and three MYB transcription factor-encoding genes, respectively (**Table S2)**. In total, 14, three, and four genes encoding MYB transcription factors were exclusively differentially expressed in responses to 6h of drought, 1h of heat, and 6h of heat treatment, respectively (**Fig.S3D**). Notably, 12 genes that encode the MYB transcription factor showed differential expression in responses to both 6h of drought and heat treatment(**Fig.S3D & Fig.5B**).

Genes encoding heat shock factors (HSF) displayed distinct expression changes in response to drought and heat stress (**Fig.6A**). In total, 19 HSF-encoding genes were differentially expressed in response to at least one of the stress treatments. 1h of drought treatment resulted in the up-regulation of only two HSF-encoding genes. However, 6h of drought treatment led to the up- and down-regulation of 11 and two HSF-encoding genes, respectively. In contrast, 1h of heat treatment elicited an early transcriptional response, with six HSF-encoding genes (including *Sobic.001G450700*) showing up-regulation in response to 1h of heat treatment, whereas only three HSF-encoding genes were up-regulated in response to the 6h of heat treatment (**Table S2**). Additionally, one, eight, four, and one HSF-encoding genes were exclusively differentially expressed in responses to 1h of drought, 6h of drought, 1h of heat, and 6h of heat treatment, respectively (**Fig.S3E**).

**Figure 6.**
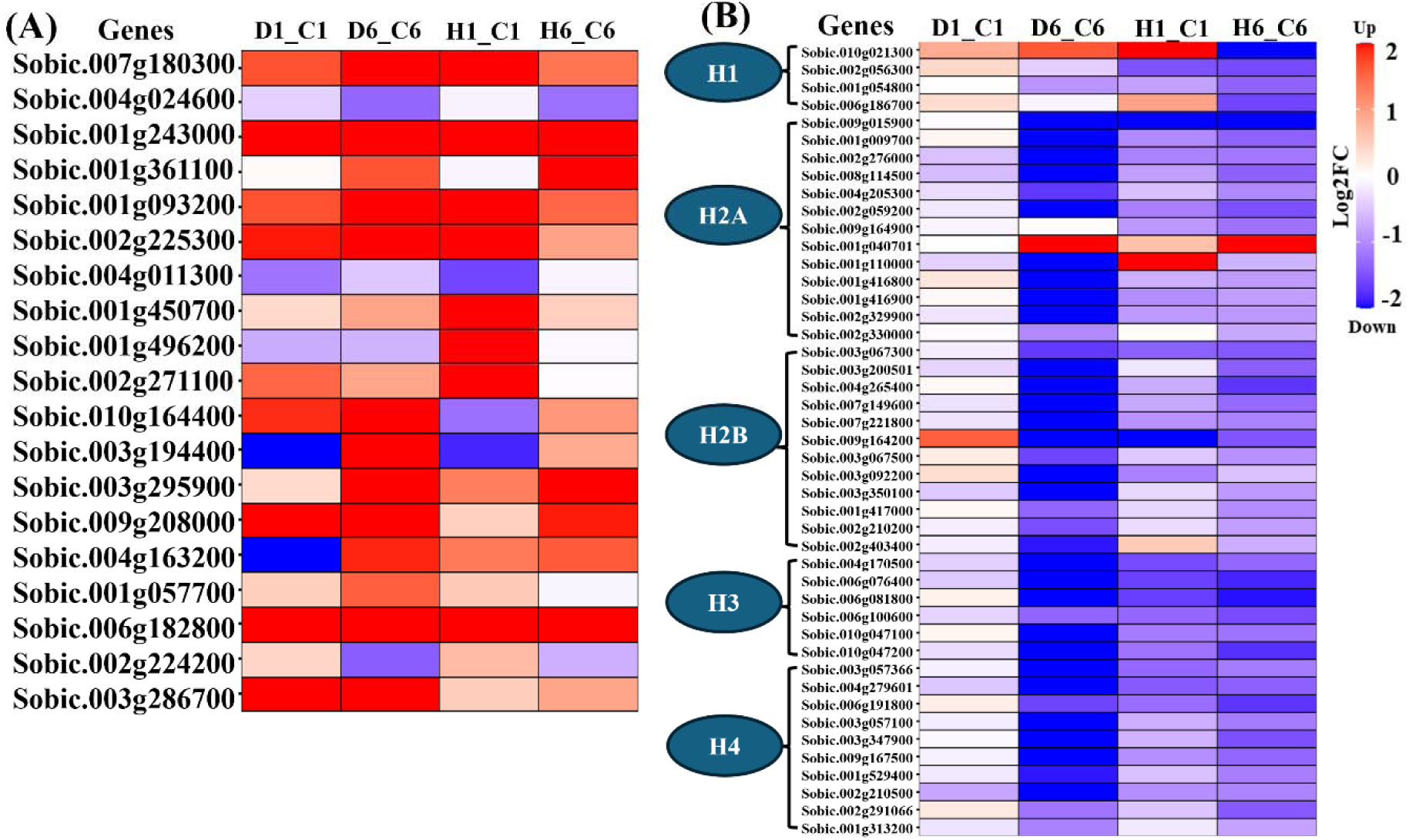
Heatmap representing DEGs encoding HSFs and histone proteins under four stress conditions. D1, H1, and C1 represent to drought, heat, and control samples at 1h. D6, H6, and C6 represent to the same samples at 6h. D1_C1, D6_C6, H1_C1, and H6_C6 represent comparisons of 1h drought-treated, 6h drought-treated, 1h heat-treated, and 6h heat-treated samples with their corresponding control samples, respectively. **(A**) represents expression patterns of genes encoding HSFs and **(B)** represents expression patterns of genes encoding histone proteins including linker histone (H1) and core histones (H2A, H2B, H3, H4) across four conditions. Fold change values represent the Logl-transformed ratio of average normalized read counts (stress/control), with red and blue indicating up- and down-regulation, respectively.

### Widespread down-regulation of histone genes in responses to drought and heat treatments

Our MapMan analysis also revealed a highly consistent downregulation of the ‘histone’ genes, specifically at the 6h drought and heat treatment, hence this set of genes was explored further (**Fig.S6**). A total of 45 genes encoding histone proteins were differentially expressed in response to at least one of the stress conditions (**Fig.S3F & Fig.6B)**. 1h of drought treatment didn’t alter the expression of genes encoding histone proteins, whereas 1h of heat treatment up- and down-regulated two and 13 genes encoding histones, respectively. However, 6h of drought and heat treatments drastically reduced the expression of genes encoding histones. In response to 6h of drought treatment, expressions of 40 histone genes were significantly altered, out of which 39 were down-regulated and just one was up-regulated. Similarly, after 6h of heat treatment, 31 genes were down-regulated, and only one histone-encoding gene was up-regulated. (**Table S2)**. A total of 11 and four genes encoding histone proteins were differentially expressed exclusively in responses to 6h of drought and 6h of heat treatment, respectively (**Fig.S3F**). Notably, 27 genes encoding histones were differentially expressed (mostly down-regulated) in responses to both 6h of drought and heat treatments as compared to respective controls. Overall, our analysis revealed a drastic reduction in expressions of genes that encode different core histone proteins (H2A, H2B, H3, and H4) in responses to 6h of drought and heat treatment (**Fig.6B**).

### Orthogonal validation of stress-responsive DEGs

We have confirmed stress-induced differential expressions of 10 DEGs identified by the genome-wide 3’-end sequencing approach in this study using an independent RNAseq data (NCBI SRA BioProject accession PRJNA523862) (**Fig.S7A to S7J**). This set of DEGs included a bZIP transcription factor-encoding gene *Sobic.010G194900*, a MYB transcription factor-encoding gene *Sobic.007G053700*, a HSF-encoding gene *Sobic.001G450700*, an ABA-responsive ABRE-binding protein-encoding gene *Sobic.004G309600*, a JA-responsive lipoxygenase-encoding gene *Sobic.001G125700*, and five genes *Sobic.002G059200*, *Sobic.004G265400*, *Sobic.006G081800*, *Sobic.003G057100*, and *Sobic.002G056300* that encode H2A, H2B, H3, H4, and H1 histone proteins, respectively. Overall, we observed a close correspondence in expression patterns revealed by the 3’-end sequencing and the RNAseq datasets, indicating robustness of the transcriptome analysis using the 3’-end sequencing approach.

## Discussion

Drought and heat stress severely impair plant growth and development, manifested by changes at the molecular, biochemical, metabolic, and physiological levels (Liu et al. 2023). Plants mount rapid and dynamic transcriptional responses to abiotic stress that can change over seconds to hours (Li et al. 2015). Our study revealed such temporal and stress-dependent transcriptional changes in response to drought and heat treatments in sorghum seedlings. In the discussion below, we focus on the evolutionary conservation of drought and heat response mechanisms across cereal species, by highlighting drought- and heat-responsive transcriptional changes observed in sorghum seedlings in our study in light of prior research findings in other cereal species, including rice, wheat, maize, *Brachypodium*, *Setaria*, and others.

In this study, we employed a genome-wide 3’-end sequencing approach, which has several advantages compared to the regular RNAseq approach. This method can be used for partially degraded RNA samples, requires lower sequencing depth, hence is less expensive, and can also be used to study alternative polyadenylation (Moll et al. 2014, De Lorenzo et al. 2017). This method is gradually gaining popularity among researchers and has been used in different plant species, including arabidopsis, rice, maize, red clover, tomato, and non-model plant species (De Lorenzo et al. 2017, Chakrabarti et al. 2018, Kremling et al. 2018, Chu et al. 2019, Ye et al. 2019, Marx et al. 2020).

### Stress-induced modulation of phytohormones and secondary metabolites, and energy allocation trade-offs in sorghum

Our transcriptomic analysis revealed that drought and heat stresses elicit both overlapping and distinct changes in the expression of genes implicated in phytohormone biosynthesis, signaling, and response, and in the accumulation of secondary metabolites (**Fig.S3, Fig.S4, and Fig.S5**). Notably, ABA-related genes exhibited strong induction, particularly in response to 6h of drought treatment, consistent with ABA’s established role as a central mediator of drought responses in plants. The up-regulation of genes encoding protein phosphatase 2C (PP2C) family members in response to the drought treatment mirrors findings in other species, where these genes are crucial for ABA signaling during water deficit. For example, in maize, *ZmPP2C15* was induced by drought and ABA, and functional studies confirmed its role in imparting drought tolerance (Pang et al. 2024). Our study also identified PP2C-encoding genes that exhibited differential expression in responses to both drought and heat treatment (**Fig.S3A & S4A**), suggesting possible roles of these genes in mediating both drought and heat responses in sorghum and in other cereal species. Similarly, Jasmonate (JA)-related signaling appears to be an integral component of sorghum’s responses to drought and heat treatments. Our data shows an induction of JA-related genes, particularly in response to 6h of drought treatment (**Fig.S3B & S4B**), suggesting that JA biosynthesis and signaling are activated as part of the sorghum’s response to drought stress. A prior study in wheat demonstrated drought-induced expression of *TaOPR2*, which encodes a flavin mononucleotide (FMN)-dependent oxidoreductase involved in the biosynthesis of jasmonic acid (Wang et al. 2016). In line with this finding, in our analysis, the *Sobic.010g084400* gene that encodes an FMN-containing oxidoreductase, displayed strong induction both in responses to 6h of drought and heat treatment, indicating a possible role of this gene in mediating responses to both drought and heat treatment in sorghum.

The heat-induced suppression of secondary metabolism after 1h, marked by significant downregulation of genes involved in lignin, phenylpropanoid, dihydroflavonols, flavonols, anthocyanin, and glucosinolate biosynthesis (**Fig. S5C**), exemplifies a physiological trade-off documented previously in cereals, including C4 crops. A prior study in pearl millet demonstrates that acute heat stress exerted a rapid downregulation of secondary metabolic pathways concurrent with the upregulation of heat shock proteins and ROS detoxification genes (Singh et al. 2024). This reflects a strategic reallocation of metabolic resources toward immediate cellular protection and protein homeostasis. Such metabolic shifts under heat stress manifest as reversible prioritization of antioxidants and molecular chaperones over energetically costly biosynthesis of specialized metabolites, underscoring an evolutionarily conserved growth-defense trade-off that maximizes plant survivability during acute abiotic stress (Zandalinas et al. 2022). Our sorghum transcriptomic data support this model, positioning secondary metabolism suppression as an early protective tool integral to cereal plants’ heat stress response mechanism.

DEGs that showed altered expression only in response to 6h of drought and that exhibited altered expression in responses to both 6h stress treatments, displayed significant enrichment of genes involved in ribosome biogenesis (**Fig.3B**). Ribosome biogenesis and the subsequent protein synthesis are some of the most energy requiring processes in a cellular system (Li et al., 2014), hence it is plausible to hypothesize that in responses to abiotic stresses (e.g., drought) sorghum plants minimize the energy expenditure by downregulating ribosome biogenesis and subsequent translation, and possibly reallocate these resources for the processes necessary for survival under an adverse condition.

### Temporal regulation of bZIP, MYB, and HSF transcription factor-encoding genes in response to drought and heat stress

In our transcriptome analysis, genes encoding several families of transcription factors, including genes that encode bZIP, MYB, and heat shock factor (HSF) transcription factors, showed distinct expression patterns in responses to drought and heat treatments. Genes encoding bZIP transcription factors exhibited significant up-regulation in response to 6h of drought treatment, but their up-regulation in response to 6h of heat treatment was much less (**Fig.5A**). The bZIP transcription factor family has long been recognized for its role in regulating ABA-responsive genes through interaction with ABA-Responsive Element (ABRE) motifs and modulating plants’ responses to abiotic stresses (Hossain et al. 2010). A prior study demonstrated that *OsbZIP71* is strongly induced by drought treatment in an ABA-dependent manner and constitutive overexpression of this gene conferred drought tolerance in rice (Liu et al. 2014). In another study, the *OsbZIP62* gene displayed induction in responses to drought and ABA treatments, and overexpression of this gene enhanced drought and oxidative stress tolerance in rice (Yang et al. 2019). Our analysis also revealed parallels to these studies, where *Sobic.010G194900*, a gene that encodes a bZIP transcription factor, showed strong up-regulation in responses to both 6h of drought and heat treatments, suggesting a possible role of this transcription factor in regulating responses to both drought and heat treatments. The induction of genes encoding bZIP transcription factors under drought stress in sorghum, particularly those linked to ABA signaling cascades, mirrors prior findings in maize, where *ZmbZIP4* and *ZmbZIP33* were induced by drought, heat, and salt stress, and were reported to function as pivotal mediators of ABA biosynthesis and abiotic stress adaptation (Ma et al. 2018, Cao et al. 2021).

The well-known transcription factor family MYB is also involved in several processes, including responses to abiotic stressors and regulation of secondary metabolism (Qi et al. 2015). Most MYB proteins found in plants are members of the R2R3-MYB subfamily. Prior research studies demonstrated their involvement in a variety of processes, such as the biosynthesis of flavonoids and isoprenoid compounds, root growth, regulation of the cell cycle, and modulation of defense mechanisms against biotic and abiotic stressors (Wu et al. 2024). A notable example is *OsMYB3R-2* from rice, which was shown to be involved in the control of the drought response (Dai et al. 2007). A R2R3-MYB transcription factor, *OSMYB55,* was found to impart heat tolerance in rice. Overexpression of *OSMYB55* in maize also led to enhanced drought and heat tolerance by activating stress-responsive genes (El-Kereamy et al. 2012, Casaretto et al. 2016). In our experiment, we also observed a notable up-regulation of R2R3-MYB genes in response to 6h of drought and heat treatments (**Fig.5B**). A recent study showed that, three MYB transcription factors MYB11, MYB12, and MYB111 form a regulatory module in conjunction with CRY1 (CRYPTOCHROME 1) and HY5 and control light-induced stomatal opening by regulating ROS homeostasis and flavonol accumulation (Chang et al. 2025). It is possible that the same CRY1-HY5-MYB regulon may play a role in modulating responses to drought and heat stress in sorghum and other cereal crops.

1h of heat treatment exerted up-regulation of expressions of genes encoding heat shock factors (HSFs) (**Fig.6A**), suggestive of a transcriptional reprogramming that allows subsequent transcriptional induction of heat shock proteins (HSPs) encoding genes, which act as molecular chaperones to help in proper protein folding and contribute significantly to managing plant responses to various abiotic stresses (von Koskull-Döring et al. 2007, Fragkostefanakis et al. 2015). A prior study in maize showed strong induction of specific HSF genes during heat stress, with possible implications in dynamic changes in histone modifications (Hou et al. 2019). The rapid upregulation of HSFs in response to 1h of heat treatment in sorghum in this study aligns with these findings and indicates a vital role of HSFs in mediating the response to heat stress in cereal crops. Among different HSF-encoding genes that were differentially expressed in response to different stress treatments in this study, *Sobic.001G450700*, encoding a putative HSF3 transcription factor, was uniquely up-regulated in response to 1h of heat treatment. A previous study in arabidopsis demonstrated that *HSF3* overexpression can induce constitutive expression of *HSP* genes even in the absence of heat stress, enhancing basal thermotolerance without detrimental developmental effects (Prändl et al. 1998). This implies that *Sobic.001G450700* may have a crucial role in mediating the initial response to heat stress in sorghum. A recent study also reported conserved transcriptional induction and regulation of *HSF* gene expression in response to early heat stress in two C4 species, maize and *Setaria viridis* (Myers et al. 2023). Our findings and these prior reports highlight a conserved role of HSFs in regulating early heat response in C4 species.

### Down-regulation of histone genes and their implications in regulating stress responses

Perhaps the most interesting change in gene expression seen in this study was the stress-associated down-regulation of core histone genes in response to 6h of drought and heat treatments (**Fig.6B**). Core histone genes, which encode H2A, H2B, H3, and H4 histone proteins, form the nucleosome core and help in packaging the DNA in the nucleus. This packaging of DNA also regulates accessibility of genomic DNA to factors that mediate DNA replication, repair, recombination, and transcription (Venkatesh and Workman 2015). A prior study reported down-regulation of core histone genes in response to drought and salinity stresses in rice (Hu and Lai 2015). In arabidopsis, infection with *Pseudomonas syringae* pv tomato DC3000 also resulted in down-regulation of core histone genes (Lewis et al. 2015). The structure of chromatin is dynamic, and canonical histones can be replaced by non-canonical histone variants, which can regulate cellular processes (Talbert and Henikoff 2010). Non-canonical histones play important regulatory roles in plants; for example, H2A.Z regulates response to elevated temperature in arabidopsis and *Brachypodium distachyon*, the ratio of H3.1/H3.3 modulates root growth in arabidopsis, and H2A.Z also modulates response to drought treatment in arabidopsis (Kumar and Wigge 2010, Boden et al. 2013, Desvoyes et al. 2014, Otero et al. 2016, Sura et al. 2017). A recent study in arabidopsis demonstrated that loss-of-function mutants of H2A.X genes (*AtHTA3* and *AtHTA5*) exhibited sensitivity to DNA damage, along with disruptions in ABA signaling, thereby linking H2A.X to both DNA repair and abiotic stress response pathways (Guo et al. 2024). It is possible that down-regulation of the expression of several core histone genes might alter the nucleosome composition within the chromatin and thus impede progression of the cell cycle during drought and heat stress in sorghum, and as a result might facilitate survival over growth in response to stress.

Beyond the core histones, the linker histone H1 is also critical for maintaining chromatin structure. Previous studies identified drought and ABA-inducible variant linker histone, H1.3, which was found to be critical for maintaining normal stomatal activity during both regular plant growth and abiotic stresses (Wei and O’Connell 1996, Rutowicz et al. 2015). We identified up-regulation of an H1.3-encoding gene in response to 1h heat and 6h drought treatments, suggesting possible roles of linker histone variants in modulating abiotic stress responses in sorghum. Altogether, our analyses show extensive down-regulation of histone genes in responses to drought and heat stress in sorghum.

## Conclusions

This study offers new insights into sorghum’s transcriptional responses to individual drought and heat stress, emphasizing both stress-induced and time-dependent expression changes. The strong induction of ABA- and JA-related genes in response to individual drought and heat treatment underscores the pivotal role of phytohormone signaling in orchestrating defense and acclimation mechanisms. Our transcriptome analysis unveiled alterations in the expression of genes that encode key transcription factors such as bZIPs, MYBs, and HSFs, which likely act as central regulators of downstream stress-responsive gene networks. Results reported in this study are based on individual drought and heat treatments. However, in the field setting, crop plants often suffer drought and heat stress concurrently. Hence, in the future, it will be worthwhile to compare the results from this study with the transcriptome-level changes in response to combined drought and heat stress in sorghum seedlings. Finally, based on the marked downregulation of core histone genes, alongside altered expression of genes encoding histone variants, we hypothesize that chromatin plasticity may be an important mechanism facilitating responses to drought and heat stress in sorghum. However, further experiments are needed to test this hypothesis. Together, these findings advance our understanding of the multilayered control sorghum employs to balance growth and survival under challenging environmental conditions. Future research may be directed towards elucidating transcriptional responses in different tissue types at different developmental stages across multiple genotypic backgrounds. Future work should prioritize functional validation of these regulatory components and explore their potential as targets for improving stress resilience in other cereal crops, thereby accelerating the translation of molecular insights into agricultural improvement.

## Author contributions (CrediT)

SB (Data curation, Formal analysis, Investigation, Methodology, Visualization, and Writing-original draft), NC (Conceptualization, Supervision, Writing- review and editing), SAG (Resources, Writing- review and editing), and MC (Conceptualization, Formal analysis, Investigation, Methodology, Funding acquisition, Project administration, Supervision, and Writing- original draft).

## Ethics declarations

### Ethics approval and consent to participate

Not applicable.

## Consent for publication

Not applicable.

## Competing interests

The authors declare no competing interests.

**Clinical trial number** No applicable

**Funding sources** This study is supported by the National Science Foundation (Award number: 2312857) (MC) and by the UTRGV Start-up grant (MC).

## Data availability

Illumina sequence data used in this article is available through NCBI-SRA BioProject PRJNA523821.

## Supporting information

Supplemental material associated with the article can be accessed in the online version of this article.

