## Supplemental Figures and Tables for "Integrative transcriptome analysis reveals distinct and common stress-responsive regulatory networks driving drought and heat responses in sorghum"

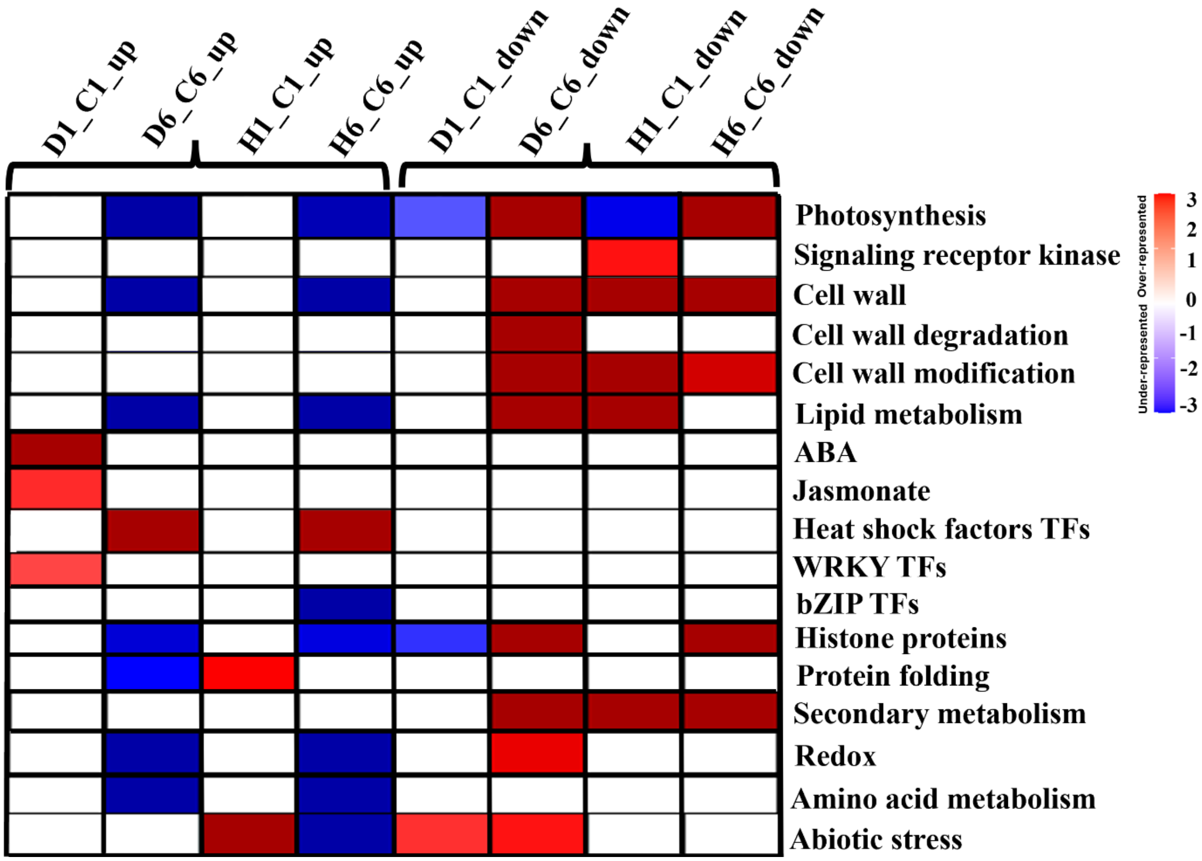


**Supplemental Figure S1. Analysis of enrichment of functional categories using PageMan among DEGs in responses to 1 and 6h of individual drought and heat treatment**. D1_C1, D6_C6, H1_C1, and H6_C6 represent comparisons of 1h drought-treated, 6h drought-treated, 1h heat-treated, and 6h heat-treated samples with their corresponding control samples, respectively. 'up' and 'down' represent up-regulated and down-regulated DEGs, respectively. The color scale represents z-transformed adjusted p-values. Red and blue indicate over- and under-represented categories, respectively. Text alongside each row denotes respective MapMan BINs.

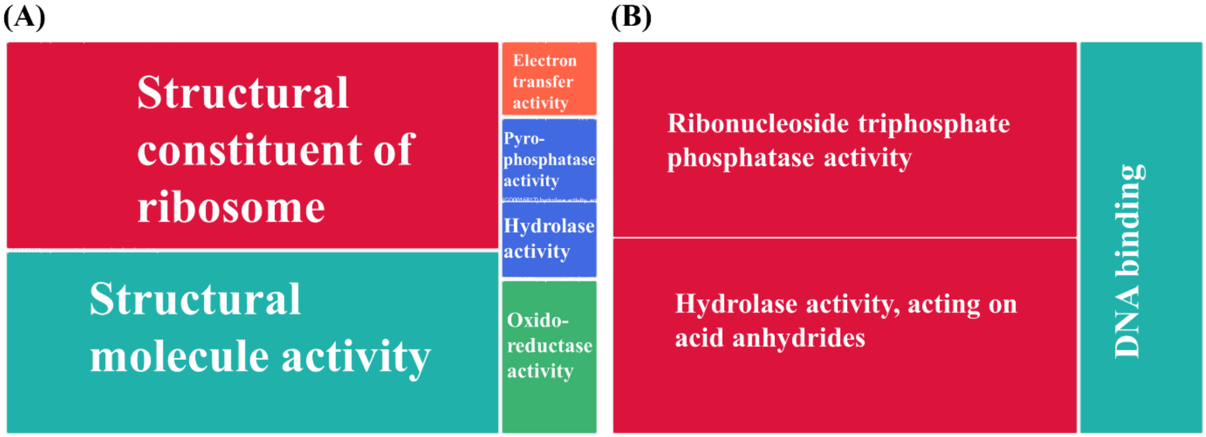


**Supplemental Figure S2. GO enrichment analysis in molecular function category for DEGs in 6h of drought and heat treatments. (A)** and **(B)** represent GO enrichment of 4136 and 2776 DEGs identified in responses to 6h of drought (D6_C6) and heat treatments (H6_C6), respectively. D6_C6 and H6_C6 represent comparisons of 6h drought-treated and 6h heat-treated samples with the control samples, respectively.

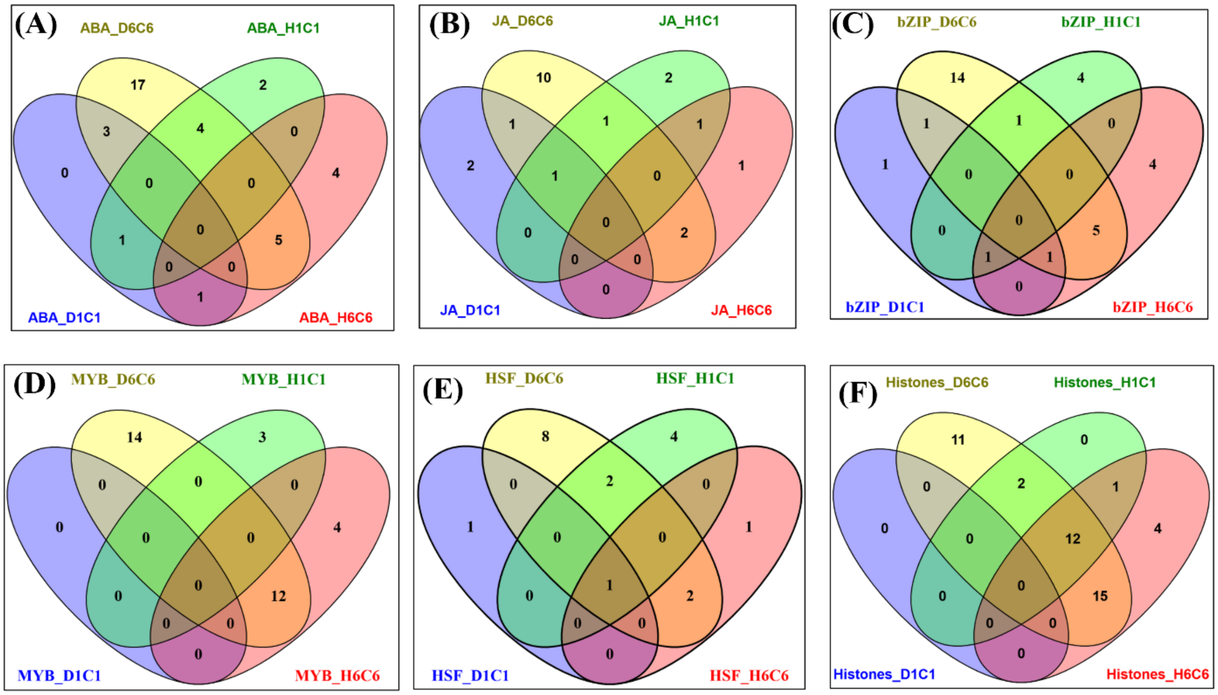


**Supplemental Figure S3. Venn diagrams showing the unique and overlapping DEGs across four stress conditions, including D1_C1, D6_C6, H1_C1, and H6_C6.** D1_C1, D6_C6, H1_C1, and H6_C6 represent comparisons of 1h drought-treated, 6h drought-treated, 1h heat-treated, and 6h heat-treated samples with their corresponding control samples, respectively. **(A)** represents DEGs implicated in ABA-related processes, **(B)** represents DEGs implicated in JA-related processes, **(C)** represents DEGs encoding bZIP transcription factors, **(D)** represents DEGs encoding MYB transcription factors, **(E)** represents DEGs encoding HSFs, and **(F)** represents DEGs encoding histone proteins.

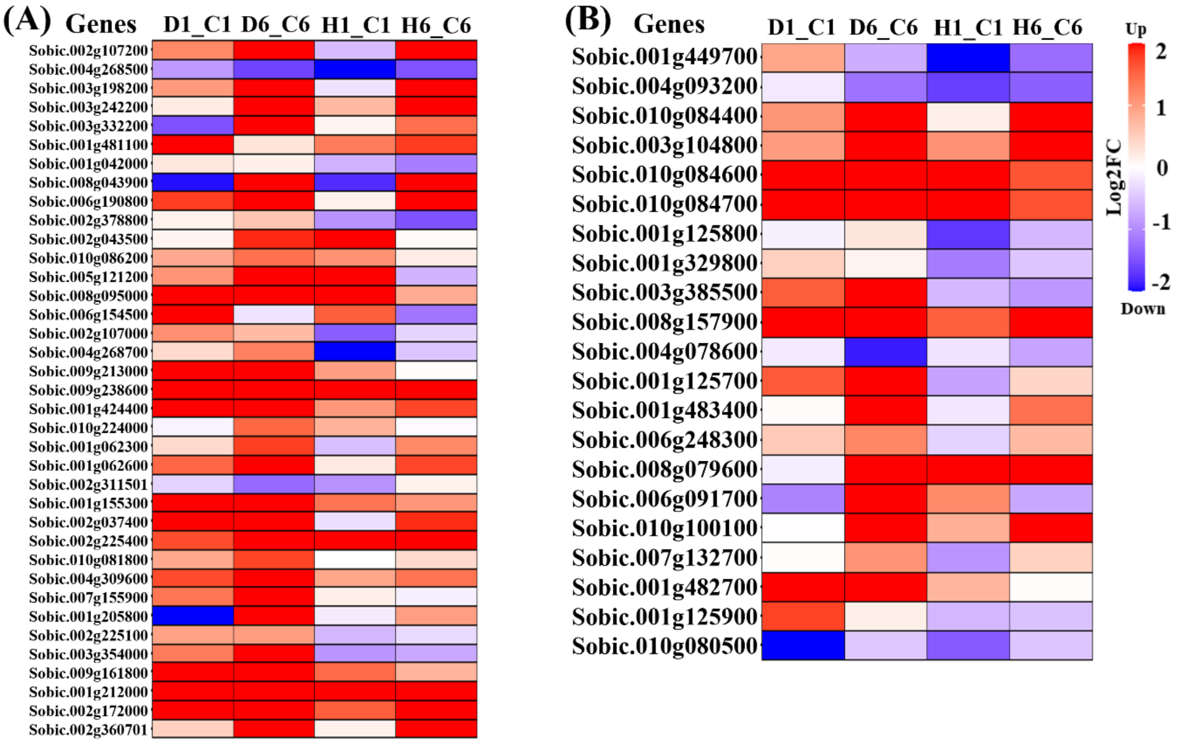


**Supplemental Figure S4: Heatmap representing DEGs related to ABA and Jasmonic Acid (JA) in responses to 1 and 6h of individual drought and heat treatment.** D1_C1, D6_C6, H1_C1, and H6_C6 represent comparisons of 1h drought-treated, 6h drought-treated, 1h heat-treated, and 6h heat-treated samples with their corresponding control samples, respectively. **(A)** and **(B)** represent expression patterns of ABA and JA related genes across four stress treatments, respectively. Gene expression values are represented by $Log_2$-transformed ratio of average normalized read counts (stress/control) across three biological replicates with red and blue indicating up- and down-regulation, respectively.

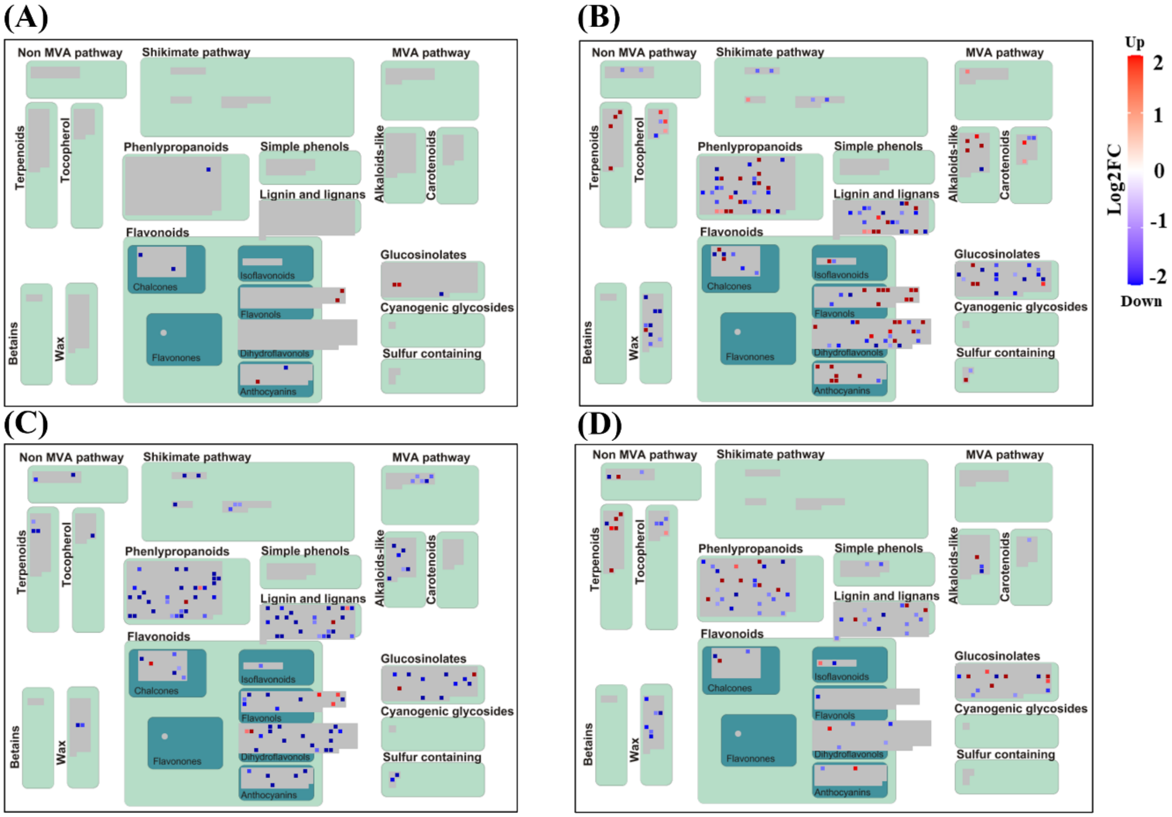


**Supplemental Figure S5. MapMan overview of DEGs involved in secondary metabolism under drought and heat treatments.** Panels show DEGs under **(A)** 1h drought (D1_C1), **(B)** 6h drought (D6_C6), **(C)** 1h heat (H1_C1), and **(D)** 6h heat (H6_C6) treatments. D1_C1, D6_C6, H1_C1, and H6_C6 represent comparisons of 1h drought-treated, 6h drought-treated, 1h heat-treated, and 6h heat-treated samples with their corresponding control samples, respectively. Gene expression values are represented as $Log_2$ Fold Change (normalized expression in stress-treated sample/normalized expression in control sample). Red and blue color indicate up- and down-regulation, respectively.

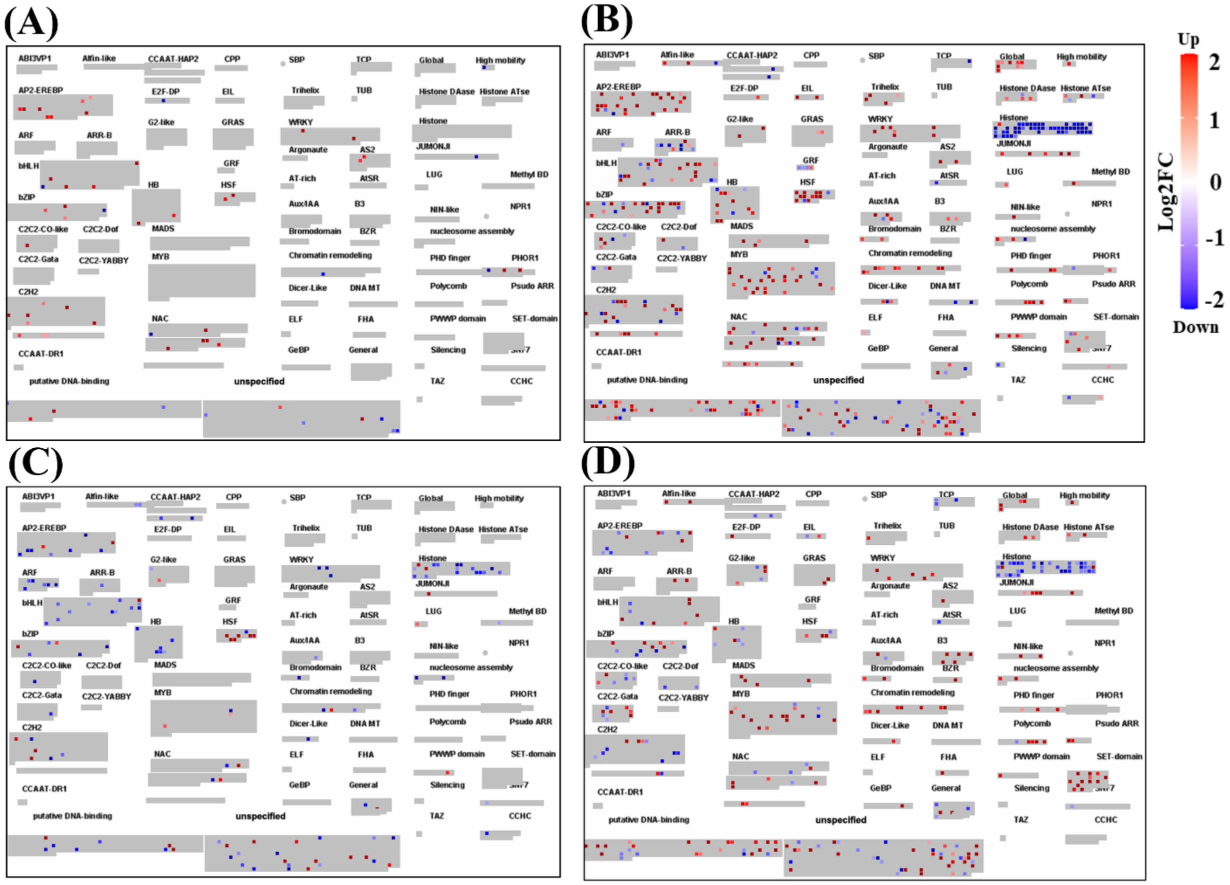


**Supplemental Figure S6: MapMan overview of DEGs involved in Transcriptional regulation under drought and heat treatments.** Panels show DEGs under **(A)** 1h drought (D1_C1), **(B)** 6h drought (D6_C6), **(C)** 1h heat (H1_C1), and **(D)** 6h heat (H6_C6) treatments. D1_C1, D6_C6, H1_C1, and H6_C6 represent comparisons of 1h drought-treated, 6h drought-treated, 1h heat-treated, and 6h heat-treated samples with their corresponding control samples, respectively. Gene expression values are represented as $Log_2$ Fold Change (normalized expression in stress-treated sample/normalized expression in control sample). Red and blue colors indicate up- and down-regulation, respectively.

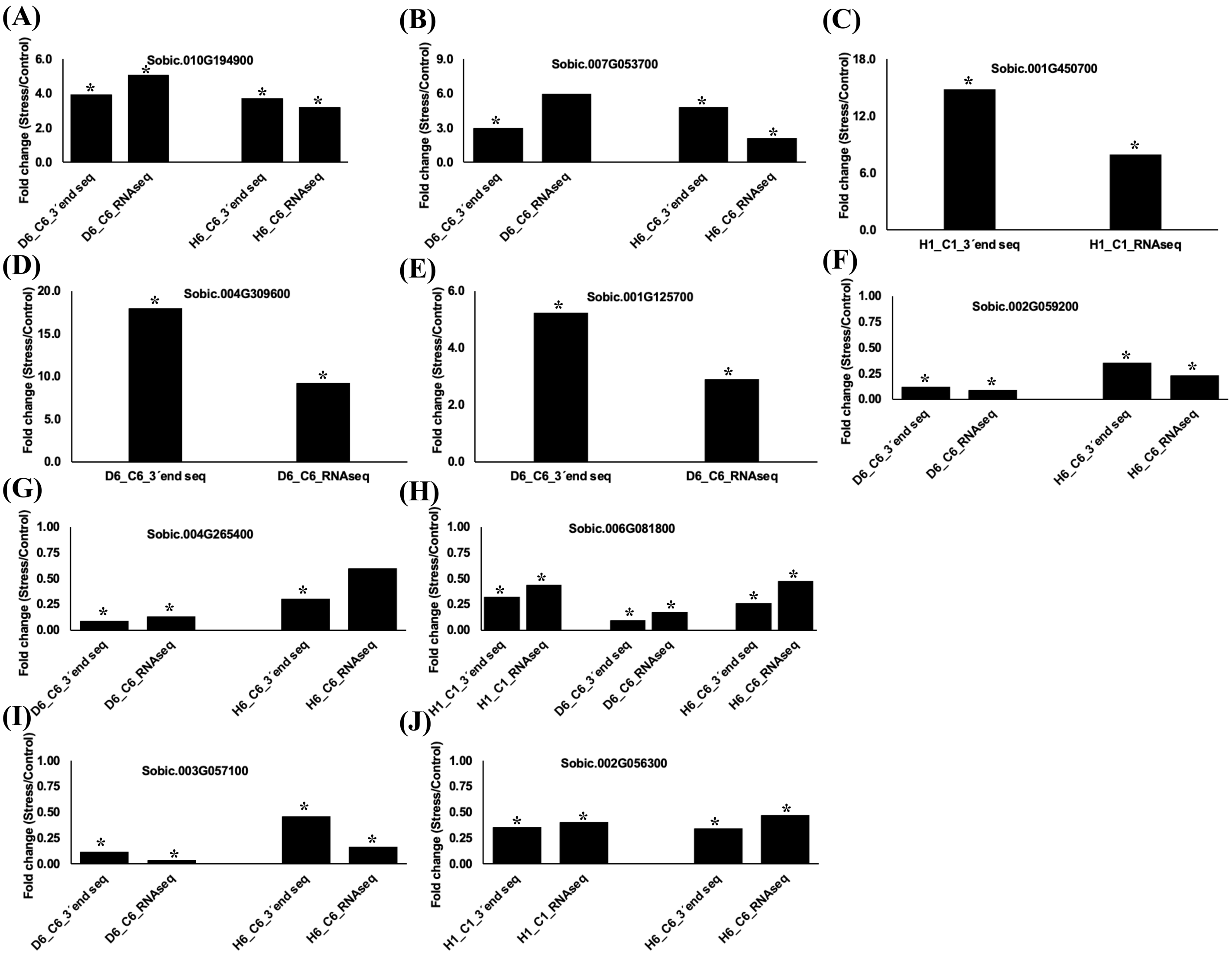


**Supplemental Figure S7. Orthogonal validation of expressions of DEGs identified in this study using the 3´-end sequencing with an independent RNAseq dataset.** D6_C6, H1_C1, and H6_C6 represent comparisons of the 6h drought-treated, 1h heat-treated, and 6h heat-treated samples with their corresponding controls, respectively. Y-axes represent fold-change values calculated by dividing average normalized read counts across three biological replicates for stress-treated samples by the corresponding values for the control samples. (*) denotes statistically significant DEGs as illustrated in the method section. 3´-end seq refers to genome-wide 3´-end sequencing.

**Supplemental Table S1: Summary of read processing and mapping statistics for sorghum samples under control, drought, and heat stress conditions.** C1, D1, and H1 represent control, drought-, and heat-treated samples after 1h, whereas C6, D6, and H6 represent the same samples after 6h. R1, R2, and R3 represent biological replicates.

| Sample | Reads after trimming | Reads retained after removing reads mapped to rRNA, mitochondrial and chloroplast genome | Reads mapped to annotated sorghum genes |
|---|---|---|---|
| **C1R1** | 12677994 | 6529427 | 5188988 |
| **C1R2** | 18427551 | 9832310 | 8365032 |
| **C1R3** | 12597040 | 6772765 | 5794896 |
| **C6R1** | 15453947 | 11039331 | 10026401 |
| **C6R2** | 13293484 | 5770110 | 4650041 |
| **C6R3** | 18194958 | 7535122 | 5864419 |
| **D1R1** | 12858216 | 5754732 | 4939369 |
| **D1R2** | 13270685 | 8977666 | 8031902 |
| **D1R3** | 14408741 | 7339515 | 6593734 |
| **D6R1** | 17816711 | 4375266 | 2509401 |
| **D6R2** | 18803919 | 3903548 | 1896753 |
| **D6R3** | 18609556 | 2903483 | 1659239 |
| **H1R1** | 13365049 | 5725343 | 4665091 |
| **H1R2** | 15375629 | 8651452 | 7179907 |
| **H1R3** | 15250549 | 7307512 | 5961967 |
| **H6R1** | 24611041 | 5775553 | 2164865 |
| **H6R2** | 23826612 | 4796266 | 1725878 |
| **H6R3** | 15786291 | 4540532 | 2285854 |

**Supplemental Table S2: Number of DEGs that are up- or down-regulated under each stress treatment for selected hormone-related pathways (ABA, JA) and transcription factor families (bZIP, MYB, HSF), as well as genes encoding histone proteins.** Values are shown for 1h and 6h time points under drought (D1_C1, D6_C6) and heat (H1_C1, H6_C6) treatments. D1_C1, D6_C6, H1_C1, and H6_C6 represent comparisons of 1h drought-, 6h drought-, 1h heat-, and 6h heat-treated samples with corresponding control samples, respectively.

| Groups of DEGs | D1C1 | | | D6C6 | | | H1C1 | | | H6C6 | | |
|---|---|---|---|---|---|---|---|---|---|---|---|---|
| | Up | Down | Total | Up | Down | Total | Up | Down | Total | Up | Down | Total |
| ABA | 5 | 0 | 5 | 27 | 2 | 29 | 5 | 2 | 7 | 7 | 3 | 10 |
| JA | 3 | 1 | 4 | 13 | 2 | 15 | 2 | 3 | 5 | 2 | 2 | 4 |
| bZIP | 3 | 1 | 4 | 17 | 5 | 22 | 2 | 4 | 6 | 6 | 5 | 11 |
| MYB | 0 | 0 | 0 | 22 | 4 | 26 | 2 | 1 | 3 | 13 | 3 | 16 |
| HSF | 2 | 0 | 2 | 11 | 2 | 13 | 6 | 1 | 7 | 3 | 1 | 4 |
| Histones | 0 | 0 | 0 | 1 | 39 | 40 | 2 | 13 | 15 | 1 | 31 | 32 |
